# ECNano: A Cost-Effective Workflow for Target Enrichment Sequencing and Accurate Variant Calling on 4,800 Clinically Significant Genes Using a Single MinION Flowcell

**DOI:** 10.1101/2021.04.05.438455

**Authors:** Amy Wing-Sze Leung, Henry Chi-Ming Leung, Chak-Lim Wong, Zhen-Xian Zheng, Wui-Wang Lui, Ho-Ming Luk, Ivan Fai-Man Lo, Ruibang Luo, Tak-Wah Lam

## Abstract

**Background:** The application of long-read sequencing using the Oxford Nanopore Technologies (ONT) MinION sequencer is getting more diverse in the medical field. Having a high sequencing error of ONT and limited throughput from a single MinION flowcell, however, limits its applicability for accurate variant detection. Medical exome sequencing (MES) targets clinically significant exon regions, allowing rapid and comprehensive screening of pathogenic variants. By applying MES with MinION sequencing, the technology can achieve a more uniform capture of the target regions, shorter turnaround time, and lower sequencing cost per sample.

**Method:** We introduced a cost-effective optimized workflow, ECNano, comprising a wet-lab protocol and bioinformatics analysis, for accurate variant detection at 4,800 clinically important genes and regions using a single MinION flowcell. The ECNano wet-lab protocol was optimized to perform long-read target enrichment and ONT library preparation to stably generate high-quality MES data with adequate coverage. The subsequent variant-calling workflow, Clair-ensemble, adopted a fast RNN-based variant caller, Clair, and was optimized for target enrichment data. To evaluate its performance and practicality, ECNano was tested on both reference DNA samples and patient samples.

**Results:** ECNano achieved deep on-target depth of coverage (DoC) at average >100x and >98% uniformity using one MinION flowcell. For accurate ONT variant calling, the generated reads sufficiently covered 98.9% of pathogenic positions listed in ClinVar, with 98.96% having at least 30x DoC. ECNano obtained an average read length of 1,000 bp. The long reads of ECNano also covered the adjacent splice sites well, with 98.5% of positions having ≥ 30x DoC. Clair-ensemble achieved >99% recall and accuracy for SNV calling. The whole workflow from wet-lab protocol to variant detection was completed within three days.

**Conclusion:** We presented ECNano, an out-of-the-box workflow comprising (1) a wet-lab protocol for ONT target enrichment sequencing and (2) a downstream variant detection workflow, Clair-ensemble. The workflow is cost-effective, with a short turnaround time for high accuracy variant calling in 4,800 clinically significant genes and regions using a single MinION flowcell. The long-read exon captured data has potential for further development, promoting the application of long-read sequencing in personalized disease treatment and risk prediction.

## Background

Screening of single genes with traditional techniques such as Sanger sequencing [1] is tedious and time-consuming, especially in heterogeneous diseases, where variants in different genes can result in similar phenotypes [2]. High-throughput sequencing allows efficient and comprehensive screening of all clinically significant genomic positions for patients with potential genetic defects. However, obtaining sufficient depth of coverage (DoC) for accurate variant calling with whole genome sequencing (WGS) is still costly for routine clinical tests [3]. Thus, target enrichment sequencing of a subset of the genome, using medical exome sequencing (MES), for example, has become a common alternative solution. MES uses customized or commercially available capture panels that target a subset of the entire exome, covering genes and positions with clinical significance. The effectiveness of exome sequencing in variant detection of rare autosomal recessive monogenic disorder [3,4] and diseases of high genetic heterogeneity [5,6,7] have been well examined. MES also requires less input DNA than WGS does [8].

Oxford Nanopore Technology (ONT) Sequencing [9,10,11] provides a cost-effective solution for long-read sequencing with minimal laboratory setup. A regular ONT MinION sequencing run generates approximately 12 to 20 Gbp data, which covers, on average, 4X to 8X of the human genome in a WGS run. ONT sequencing involves real-time generation of molecular signatures while the nucleotide polymer passes through a biological CsgG protein pore [11], which can be used for both DNA and RNA sequencing, as well as nucleotide modification detection. The simultaneous base-calling while sequencing significantly shortens the data-processing time. The technology, therefore, has diverse potential applications in genomic medicine [12].

Owing to the high sequencing error rate of ONT, it remains challenging to use a single MinION WGS run to achieve high-quality variant detection. Target enrichment resolved the issue by improving the DoC in clinically important regions. A recent development of ONT sequencing incorporated the use of CRISPR/Cas9, which can effectively capture and enrich large genomic fragments for single-nucleotide variants (SNV) and structural variant detection. However, CRISPR/Cas9 targeted ONT sequencing can be performed on only a small scale, tested on 10 loci [13]. The sequencing cost, therefore, is still high in routine clinical applications targeting over 1,000 genes. In addition to the CRISPR/Cas9 enrichment protocol, ONT has developed an amplicon sequence capture protocol that can be applied to exome sequencing. The protocol can be performed with an average DoC of about 30x on whole-exome sequencing [14], which is insufficient for high-quality variant calling, especially for positions with <30x DoC. Further optimization is needed to increase the average DoC.

A couple of existing bioinformatics tools are available for preprocessing and variant detection using ONT data. The recommended data processing pipeline on the ONT proprietary analysis platform EPI2ME (https://epi2me.nanoporetech.com/) provides quality control of data based on the alignment result. The workflow is incomplete without further downstream analysis. The existing third-generation sequencing (TGS) variant calling tools, such as Medaka (https://nanoporetech.github.io/medaka/index.html), LongShot [15],and Clair [16], are designed and trained mainly for WGS data, assuming a relatively even DoC in the sampled regions.

However, owing to the variation of capturing and PCR efficiency in different targeted regions, the average DoC among the captured blocks fluctuates across the genome. Some of these variant callers generate consensus sequences and perform haplotype phasing [17,18] for error correction in their workflow, which significantly increases the runtime and lowers the sensitivity in high DoC regions. While the availability of high-depth models could improve the performance of callers at high DoC positions, this is not as sensitive as some low-depth models at positions with 100X or below.

In this study, we developed a workflow, ECNano, for accurate variant calling of MES of 4,800 clinically significant genes using a single ONT MinION flowcell. The workflow comprises (1) a wet-lab protocol for the target enrichment ONT sequencing and (2) a bioinformatics pipeline, Clair-ensemble, for subsequent variant calling. The ECNano wet-lab protocol was designed to work with the solution-based target enrichment Agilent SureSelect Focused Exome panel, which strikes a balance between panel size and obtaining a sufficient DoC for high-quality variant calling in one sequencing run. Since the average exon size is about 164 bp [19,20], our workflow targets an average fragment size of 1,000 bp, which uniformly covers the targeted exome region, as well as the close-by splice sites. Alignment quality, and therefore variant-calling accuracy, were significantly improved with long reads. Another advantage of longer reads is obtaining more even coverage in positions with nearby homopolymers and small repeats, which is often excluded in the design of the capture probes [21]. The bioinformatics pipeline Clair-ensemble adopts the fast ONT variant caller Clair [16] for variant calling. Clair can achieve more accurate and refined variant classification, including the multiple bi-allelic variants, as well as the long INDEL variants. To improve the performance of variant calling with amplicon data, Clair-ensemble applies a by-positional subsampling strategy for high DoC positions and provides the ensemble of the results of Clair using multiple models. To ensure stable performance and reproducibility, the ECNano workflow was tested with both standard reference DNA and patient samples. Good-quality, high-coverage long-read data were generated for high-accuracy single-nucleotide polymorphism (SNP) calling. The precision of INDEL calling with ONT data was also significantly improved, and was benchmarked against other variant callers. The whole workflow was completed within three days. The application is therefore suitable for urgent genetic testing. Further development of the workflow is promising, extending towards effective variant phasing and trio analysis. This work is also significant in promoting the application of long-read sequencing in personalized disease treatment and risk prediction.

## Method

### Overview

The ECNano workflow, comprising a wet-lab protocol and bioinformatics analysis, can be completed within three days, including (1) target capture, (2) target enrichment, (3) ONT library preparation, (4) MinION sequencing, (5) data preprocessing, and (6) variant calling. Our protocol is optimized for the use of SureSelect^XT^ Focused Exome (Agilent, Santa Clara, CA, USA), which targets approximately 17 Mbp positions. Other custom panels of similar target size are also expected to be applicable. The summary workflow within the three-day time-frame is illustrated in Fig. 1.

**Fig. 1.**
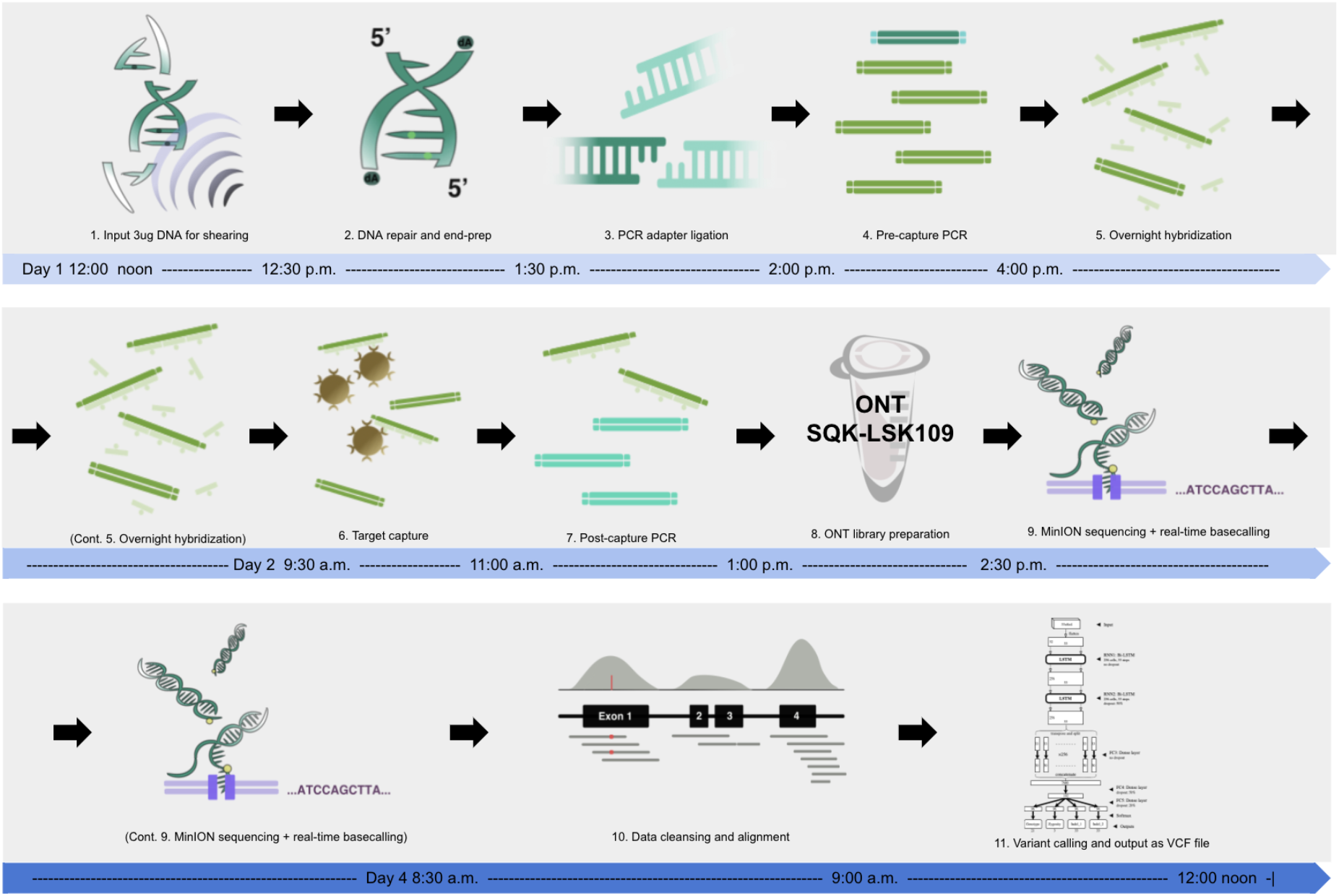
Overview of the ECNano workflow diagram. The workflow comprises both wet-lab and bioinformatics components and was completed within three days (Day 1 noon to Day 4 noon).

### Wet-lab protocol of ECNano

The patient DNA samples used in this study were extracted from EDTA blood samples. The automated DNA extraction was performed using the Promega Maxwell® RSC Instrument (Promega, Madison, WA, USA) with the Promega Maxwell® RSC Blood DNA Kit. The pure standard DNA sample HG001 and HG002 were purchased from Coriell Cell repositories. In our protocol, the following experimental procedures and parameters were optimized on the basis of the ONT sequence capture protocol using the SQK-LSK109 ligation kit for the SureSelect^XT^ Focused Exome capture panel:

1. *Target fragment length:* To obtain a stable protocol that gives sufficient throughput using a single MinION flowcell for variant calling, we tested the workflow on a target fragment length of 300 bp, 700 bp, 1,000 bp, and 2,000 bp. The sequencing throughput is optimal and was therefore fixed at the fragment length of 1,000 bp. Based on the target fragment size, a DNA purification procedure after each reaction was performed using 0.6X Agencourt AMPure XP beads and washed with 80% ethanol to retain the most DNA fragments at about 1,000 bp.
2. *DNA shearing:* We tested the feasibility of using Covaris microTUBE AFA Fiber Snap-Cap with the M220 Focused-ultrasonicator as a better alternative for DNA shearing compared to DNA fragmentase. The input DNA amount was tested with approximately 3.5ug, which provided spare DNA for the construction of extra libraries if needed. It is also possible to use less DNA as input (2–3 μg) for downstream processing if materials are limited. The DNA volume was adjusted to 130 µl Tris-HCl (pH 8.0) and was sheared into 1,000 bp fragments using Covaris microTUBE (Covaris, Inc., Weburn, MA, USA) with the following settings: 20 °C, 50 W peak incident Power, 2% Duty Factor, 200 cycles per burst, and 20-second treatment time. Optional validation steps with QIAxcel Advanced System (Qiagen, Hilden, Germany) or a run on 1% TBE-agarose gel electrophoresis can be done to check the quality of shearing and AMPure XP beads clean-up.
3. *End-repair reaction:* There were significantly more DNA fragments and thus more free-ends to be repaired in a solution of 1,000 bp DNA compared with those of ultra-high molecular weight DNA (i.e. >10kbp). For the DNA repair and end-prep step, the reaction was incubated at 20 °C for 10 mins and subsequently at 65 °C for 10 mins on a thermocycler, instead of 5 mins each in the ONT sequence capture protocol. The increase in incubation time improved the yield of end-repaired DNA, with larger amounts of fragments.
4. *Pre- and post-capture PCR reaction:* We tested the performance of Tks Gflex™ DNA Polymerase (Takara Bio Inc., Japan) and LongAmp™ Taq 2× Master Mix (New England Biolabs, Ipswich, MA, USA) for 1000 bp fragment amplification. The use of LongAmp™ Taq with PRM primers included in the ONT extension kit EXP-PCA001 (ONT) performed better. The PCR condition was optimized to generate enough PCR product for downstream library preparation, thus avoiding the need for further secondary PCR amplification. The settings of the PCR cycles were as follows: (1) initial denaturation 95 °C for 3 mins; (2) 14 cycles for pre-capture PCR and 17 cycles for post-capture PCR of denaturation at 98 °C for 20 secs, followed by annealing at 62 °C for 15 secs and extension at 65 °C for 3 mins; and (3) the final extension at 65 °C for another 3 mins.
5. *Target enrichment*: The ECNano protocol uses the Agilent SureSelect^XT^ Focused Exome (Cat no. 5190-7787) capture panel with SureSelect TE Reagent kit, PTN (Cat no. G9605A) for target capture and enrichment. The target enrichment kit was originally designed for Ion Proton sequencing, which is more suitable for our target fragment length than those designed for Illumina sequencing. Most of the steps were conducted following the manufacturer’s protocol for probe library size over 3 Mb. Note that during hybridization and capture, a lower rotation speed of 1,400 rpm was used during shaking incubation at room temperature on a 96-well plate mixer for long biotinylated RNA–DNA complexes to bind better on the Streptavidin magnetic beads. After the hybridization and enrichment steps, the end product yielded approximately 1 µg purified amplicons for a standard ONT SQK-LSK109 library preparation.
6. Sequencing proceeded for 24 to 48 hours with a MinION flowcell. Signals were recorded in fast5 files by MinKNOW (ONT), ready for base-calling and downstream analyses.

After we completed our experiments, ONT released an upgraded ligation kit, SQK-LSK110. The new kit is advertised as an in-place replacement for SQK-LSK109. Our protocol can use the new kit without any changes, and we expect to see a similar or better read length distribution and on-target rate using the new kit.

### Bioinformatics workflow of ECNano

The bioinformatics workflow and test data are available at GitHub (https://github.com/HKU-BAL/ECNano). We benchmarked the performance using existing ONT variant callers, including LongShot, Medaka, and Clair. We tested, but excluded, PEPPER (https://github.com/kishwarshafin/pepper) from our benchmark because of its low running speed and lack of model support for exome sequencing. While Clair performs the best in terms of sensitivity and precision, the calling time is significantly shorter than the others. We therefore further optimized a variant detection workflow, Clair-ensemble, for the ECNano exome capture data. The bioinformatics workflow could also be applied to other ONT amplicon data with uneven depth distribution among the targeted regions. The major adaptations are as follows:

(7) *Data preprocessing*: The fast5 signals generated were real-time base-called using ONT basecaller Guppy v.3.4.4 using HAC config during sequencing. In addition, homopolymer correction and a minimum q-score of 3 were set for base-calling. The adapters in reads were trimmed using Porechop v0.2.1 (available at https://github.com/rrwick/Porechop). Reads with middle adapter identified were removed or split, based on a user-defined setting to avoid possible chimeric reads that might cause false positives during alignment. Reads were aligned to human reference genome hg38 (GRCh38) using minimap2 with the ONT genomic read alignment setting [21].
(8) *By-position resampling*: To ensure only high confidence alignment was retained for variant calling, only primary alignments with mapping quality of 60 or above were retained. Instead of retraining the model using capture data with different sequencing depths, we proposed a more flexible method of resampling the sequencing data into multiple datasets for each high-depth region. The resampling function is a customized partitioning method embedded in Clair. During the initial data preprocessing steps, the user (1) can specify the maximum depth per position within the target BED region, (2) can specify the maximum number of partitions to be subset for positions above the specified maximum depth, and (3) can apply optional base quality filtering at a subsampled position before variant calling when resampling is applied. The partitioning is mostly non-overlapping. If the last partition has a smaller number of alignments than the specified depth, extra alignments can be resampled from the total reads until the required depth is reached. To preserve the sensitivity of variant calling in low-depth positions (e.g. below 10x), resampling and the base-calling filter are not applied if the depth is below the maximum depth cut-off value, which can therefore minimize the information lost, especially in low-depth regions before variant calling. In our benchmarks, we set the maximum depth at 100x, with a maximum of 5 partitions in positions that required down-sampling, and set the base quality cut-off at q-score 5.
(9) *Ensemble variant calling*: Variant calling on each partition of candidate positions was performed using Clair [16], with four models trained on different combinations of ONT WGS reference data. The details of each model used are listed in Table 1. Clair performed individual classification tasks on genotype, zygosity, INDEL length per haplotype, and output as a probability. Variant calling was performed in each of the candidate variant positions within the MES bed region using these models to obtain the probability of each classification task. To ensemble the classification results of the four models, the probability of each task per called position was averaged across partitions and models used to provide the most robust probability calculation. The averaged probabilities were used for post-processing of Clair to decide on the variant calling output.

**Table 1.** Information on the four variant calling models of Clair used in Clair-ensemble (CE) for variant detection of ECNano data. The models were trained with combinations of different ONT WGS datasets (Supplementary files in Ref. [16]). * Normal depth models were trained with datasets of maximum 168x DoC, and the model with mixed high depth data included a dataset of maximum 578x DoC.

| Model | Training set | Specifications* |
| --- | --- | --- |
| CE1-3-4 | hg001 + hg003 + hg004 ONT data | Normal depth WGS model |
| CE1-2-2HD-3-4 | hg001 + hg002 + hg002 very high depth (up to 500x) + hg003 + hg004 ONT data | Very high depth and normal depth WGS mixed model |
| CE1-2 | hg001 + hg002 ONT data | Normal depth WGS model |
| CE1-2-3-4 | hg001 + hg002 + hg003 + hg004 ONT data | Normal depth WGS model |

We tested the alignment down-sampling performance using VariantBAM [22] with maximum coverage option (i.e., m option) and SAMtools random subsampling (random seed set using -s option). For variant calling, we evaluated the performance on ECNano target enriched ONT data using LongShot, Medaka, and Clair. The benchmarking was done following the standard described in detail in Clair. The output was generated in standard VCF format, which can be used as input for other downstream processing, such as variant phasing.

## Results

### Reference DNA and clinical samples for performance evaluation

The wet-lab protocol of ECNano was tested using the standard HG001 and HG002 DNA samples, which are commercially available, ensuring that the performance assessment of the wet-lab protocol was unrelated to the quality of input DNA. In addition, these two datasets are among the best-annotated human references, which allows precise variant calling evaluation of Clair-ensemble using ECNano data. To assess the reproducibility of ECNano workflow in actual clinical practice, ECNano was tested on three in-house clinical samples that require detection of different variant types and genotypes. The patients were diagnosed, and disease-causing pathogenic variants of the samples were previously identified using the next-generation sequencing approach. Our workflow therefore performed well using standard reference samples and was sensitive to pathogenic variant detection using clinical specimens.

### Stable, high-quality performance in long-read target capturing for MinION sequencing

Among the various target enrichment methods, using RNA or DNA biotinylated probes for hybridization-based target capture is stably applied with short-read sequencing for genetic screening in clinical genetics laboratories. The method is standardized and robust, and therefore highly reliable. ECNano therefore integrates hybridization-based medical exome capture with MinION sequencing, which can be easily incorporated into existing clinical practice as a stable enhancement of the genetic screening workflow. To ensure the workflow was very steady, we tested the balance between the capture efficiency of Agilent SureSelect Focused Exome probes and fragment length. The workflow was optimized to capture input DNA fragments of approximately 1,000 bp. After MinION sequencing, the N50 of the captured reads was 1,378 bp, and the average read length was 968 bp after minimum quality filtering and adapter trimming (Fig. 2).

**Fig. 2.**
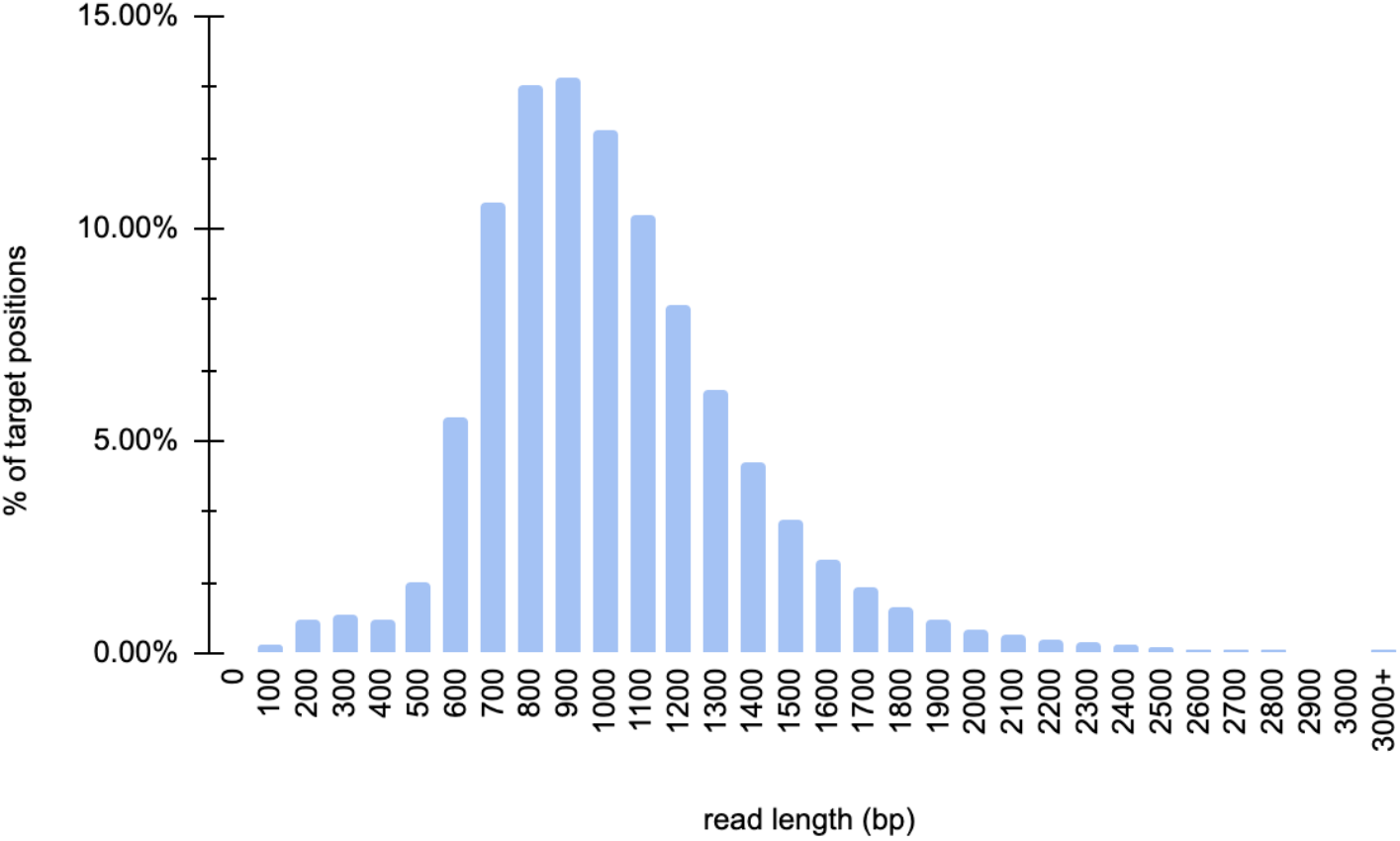
Read length distribution of an ECNano target captured HG001 library, sequenced using an ONT MinION sequencer with a single flowcell with 10Gbp throughput.

Per-base accuracy is often constrained by sequencing technology. Low-quality reads with a mean Phred score below 3 were removed during base-calling. The average read Phred score in the tested sequencing library was Q11, and the best was Q24. Over 70.2% of the reads achieved Q10. Although the highly accurate base-calling model of base-caller Guppy was used, the error rate of ONT reads was still high, so conducting accurate variant calling is still challenging if the DoC is not high enough. The DoC per position deviated from the sequencing throughput. Depending on the quality and number of active pores available in the flowcell used, each sequencing run is expected to yield minimum 10Gbp throughput. Even with a sequencing run of only 10 Gbp throughput, ECNano achieved DoC at target positions of 133x, on average. About 98% of the target positions listed in the panel were covered by at least 30 reads, about 87% of the positions had 60x DoC, and about 55% of the positions had 100x DoC (Fig. 3). With long ONT reads, over 96% of reads could be uniquely mapped to the reference genome for variant calling. Out of the 72,417 target regions within the MES bed, on average, only 17 regions did not have any read covered. Most of these regions had over 80% GC content, so they might be difficult to capture.

**Fig. 3.**
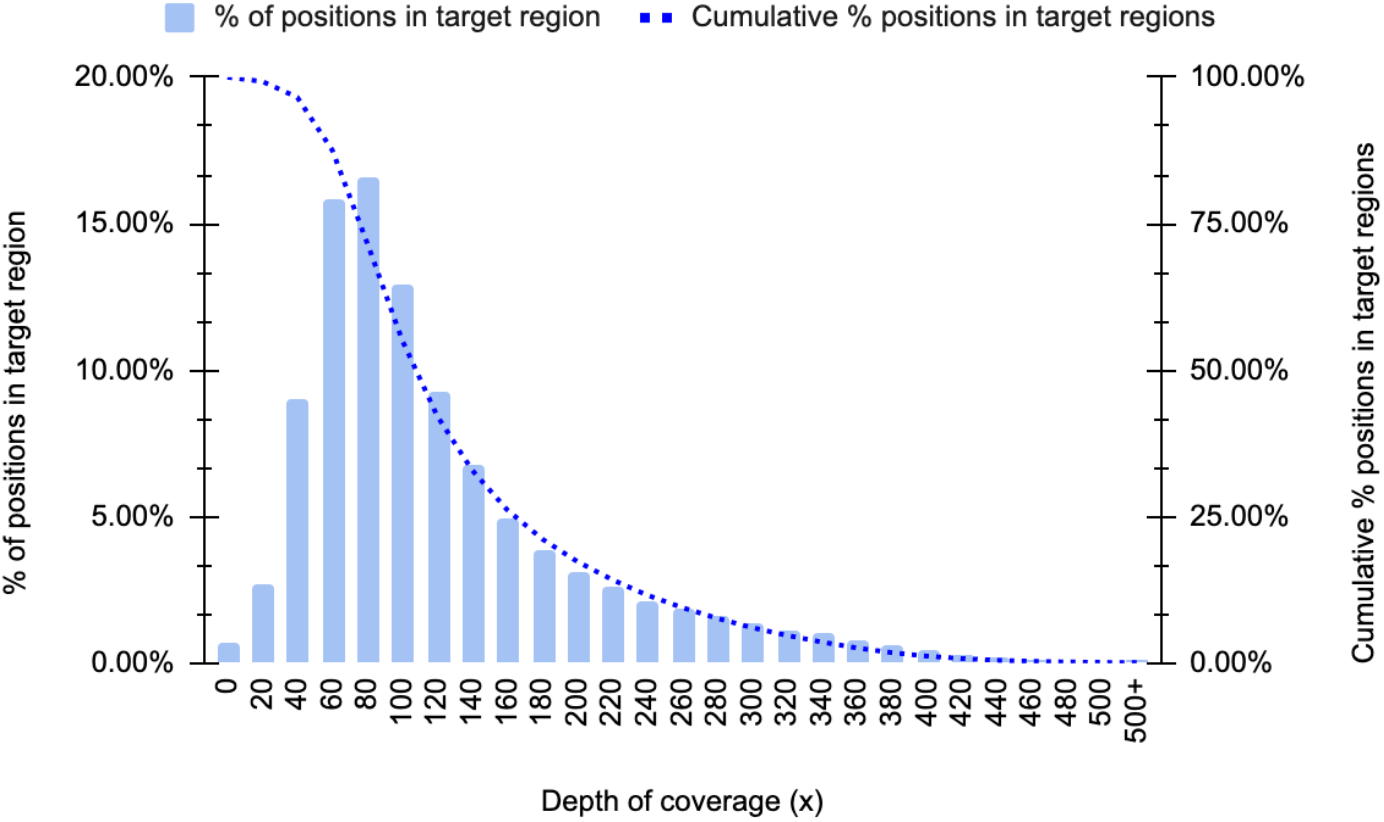
Depth distribution of the ECNano target positions in a sequencing run using standard reference HG001 samples. Positions with 30X coverage or above are considered to be highly confident for variant calling.

The margins of the target exon were uniformly covered by long reads of ECNano. The uniformity of the sequencing data was over 98% in the target regions. For adjacent exons with a short intron between them, our data showed a broader covered region, and the intron positions were also covered (Fig. 4). This allows ECNano data to have high discovery potential for novel pathogenic variant detection in a broader range around these medically relevant positions. Although there were still considerable deviations in the DoC in the captured regions owing to variation in capture efficiency and PCR bias, in-depth normalization was well performed by Clair-ensemble to achieve precise variant calling.

**Fig. 4.**
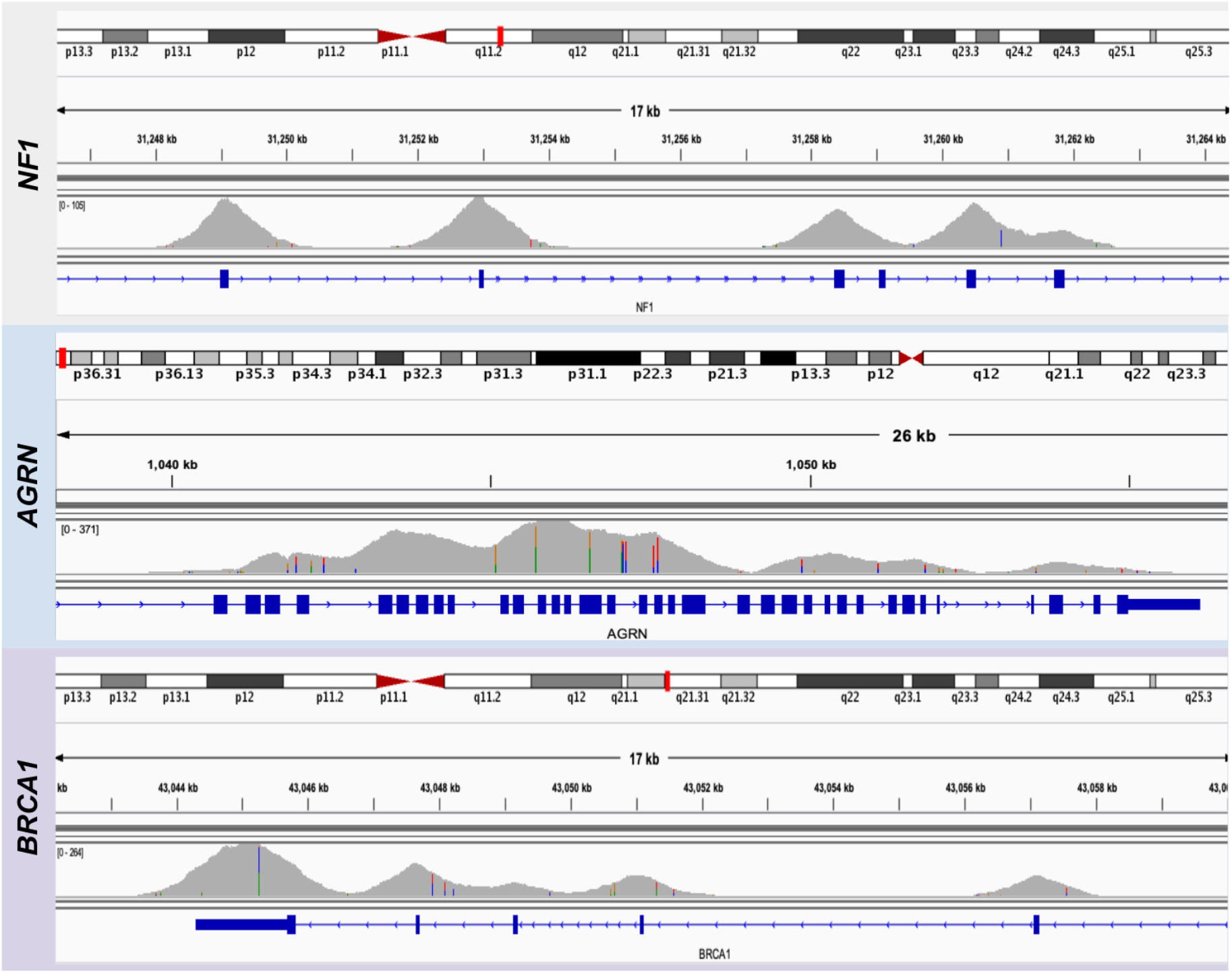
Sample depth distribution comparison of target regions in three genes: (upper: *NF1*; middle: *AGRN*; bottom: *BRCA1*) in HG001 with ECNano target enrichment protocol.

### Sufficient DoC is guaranteed at known pathogenic variant positions and nearby splice sites for accurate variant calling

Known pathogenic variant positions and nearby splice sites are most relevant to medical diagnosis. Since most of the recorded pathogenic missense variants were found in protein-coding sequences [23,24], screening variants in exome regions is sufficient for predicting a large proportion of genetic diseases. It is essential to maintain sufficient DoC at these positions to guarantee that the variant detection is highly sensitive. To evaluate the performance of target enrichment at the known pathogenic variant positions, we examined 23,866 positions in the targeted region of ECNano, which are listed in the ClinVar database as pathogenic using one of the HG001 sequencing runs. Over 98.9% of these positions achieved above 30x DoC, which allows highly confident variant calling at these critical positions.

Since the average size of an exon is 164 bp [19, 20], a library with a read length of over 1,000 bp can cover the entire exome region, as well as potential variants close to the splice sites. In total, 2,114,539 positions of potential splicing donors and receptors adjacent to the target region were extracted from the intropolis database [25]. The referenced study annotated potential donor and acceptor splice sites on the reference genome using 21,000 NGS transcriptome data. Over 90% of these positions (± 10bp included) had at least 60x DoC, and 98.5% had at least 30x DoC. This coverage allows accurate identification of canonical pathogenic splice sites and their nearby variants, which are known to cause some monogenic disease and developmental disorders [26,27].

### Accurate variant calling and performance enhancement with Clair-ensemble in target enrichment datasets

To ensure consistent and fair evaluation of variant-calling performance, Clair-ensemble was evaluated using the two sets of standard HG001 and one set of HG002 ECNano target-enrichment data against the known variants in the GIAB truth set (Table in Additional file 1). Following the best practice in Luo et al. [16], positions in high confidence regions defined by GIAB were used for benchmarking. In total, 14,976,390 positions were included in the benchmarking, which was approximately 84% of the Agilent BED positions. We benchmarked the performance of Clair-ensemble against the original Clair, LongShot, and Medaka. They are listed by precision and efficiency in Table 1. In addition to different variant callers, another downsampling method using VariantBam before variant calling was tested.

All the evaluated tools, including the original Clair, LongShot and Medaka, are capable of SNP detection. With our high DoC ECNano data, the SNP calling accuracy of these tools does not significantly deviate from each other. However, the sensitivity is inferior with Medaka, in addition to its lengthy calling time, possibly because of the use of consensus calling. The performance of Indel prediction with Medaka was also dissatisfactory. The performance of LongShot and the original Clair with low-DoC WGS models was similar in SNP calling. However, LongShot does not provide a function for Indel detection. Although we still included LongShot in our benchmarking, it is not recommended in the ECNano workflow for clinical context since it is functionally incomplete. Among all the benchmarked tools, Clair-ensemble achieved the best precision and sensitivity, achieving an F1-score in the overall calling results of over 98%. Both the built-in down-sample option and VariantBam were tested in Clair-ensemble. The calling results were similar with both resampling methods, with the built-in resampling method having slightly higher precision and overall performance. However, the use of the built-in method is preferred since no extra time or disk storage is required. It is expected that with runs of even higher throughput, the built-in resampling method will be much more effective than using VariantBam because of the increase in DoC variation with regions. The performance of the ECNano bioinformatics workflow was significantly better than the original Clair, especially for INDEL calling (Fig. 5; Supplementary Table 1). Accurate INDEL prediction with ONT data is difficult owing to random base shifts during base-calling. Although the sensitivity remained at a similar level, the ensemble method improved the precision in INDEL calling, which is preferred in clinical applications.

**Fig. 5.**
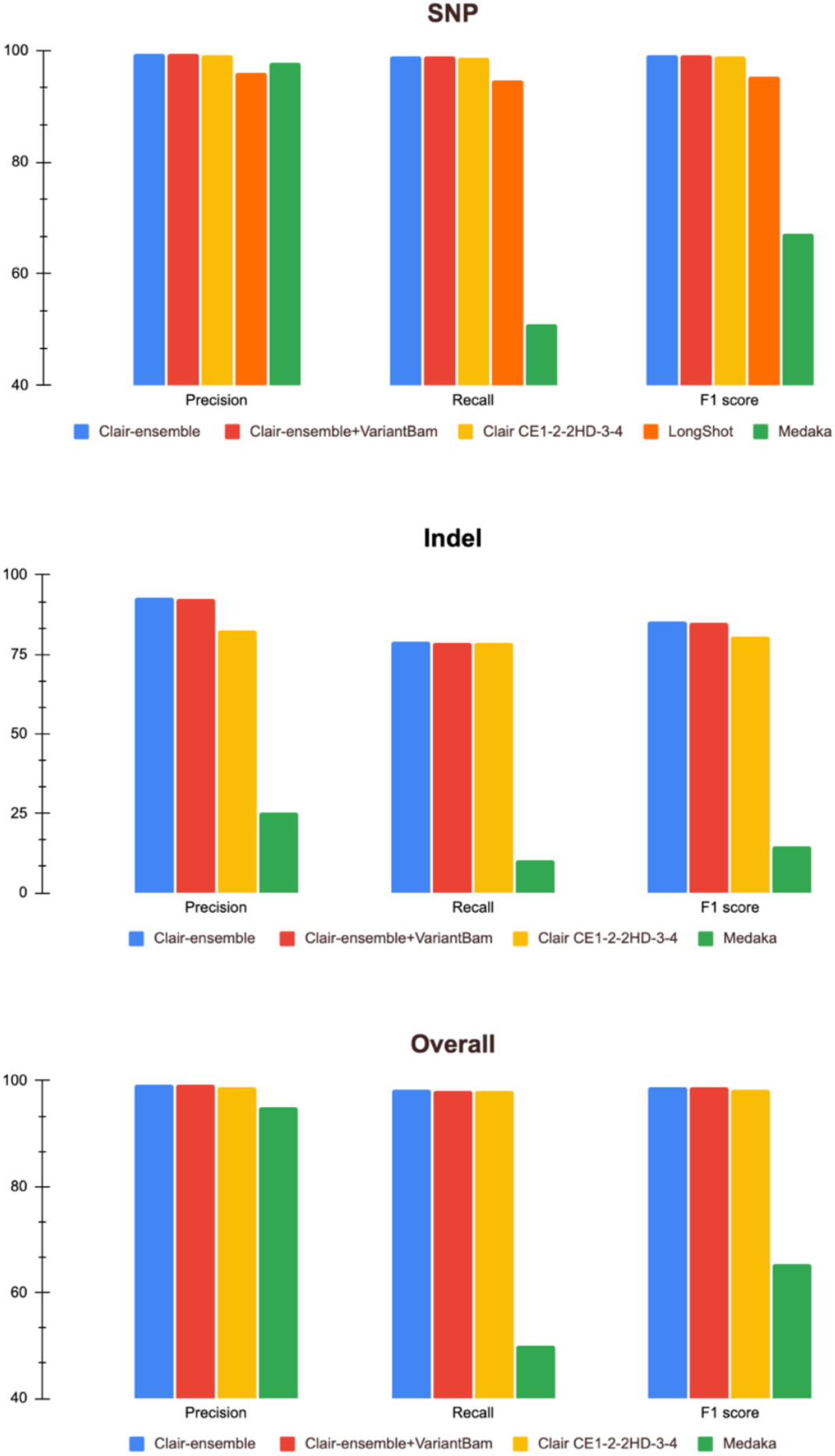
Performance of Clair-ensemble against other existing ONT variant callers at target positions using an ECNano HG001 dataset. The performance was evaluated in terms of overall (top), SNP (middle), INDEL (bottom). Clair-ensemble was performed with both the built-in resampling method and down-sampling with VariantBam. Other tools evaluated included the original Clair with different ONT models (model details described in Table 1); and LongShot for SNP calling.

The performance of neural networks is biased towards the properties of its training dataset. These variances among callers or models are reduced with the ensemble method, achieving a more stable performance across datasets. Including multiple outputs for an ensemble, however, is time-consuming if the caller runtime is long. In a comparison of the calling speed, a single Medaka run on ECNano data took days, while the original Clair required only hours. The short processing time of Clair allows an ensemble of multiple calling results without requiring too much time.

Another important feature of Clair-ensemble is the per positional resampling as the preprocessing function alongside the ensemble. The resampling function is implemented in a Clair preprocessing step, which ensures the calling DoC does not exceed the maximal allowed depth in the ultra-high DoC region, while no resampling is performed in regions with optimal or low DoC. Compared with the use of other global downsampling methods, such as VariantBam and Samtools, this is particularly effective in avoiding over-down-sampling in low DoC regions. A huge variation in DoC is commonly observed in other target enrichment data and is also highly applicable for processing with Clair-ensemble. The resampling process in Clair-ensemble also preferentially retains higher quality bases for variant calling, and therefore further improves the precision of variant calling.

### Practical application of ECNano on real patient samples

Since the whole wet-lab protocol optimization and variant-calling performance evaluation was completed with standard DNA samples, we also applied the complete workflow using three patient DNA samples to ensure that ECNano is practical in actual clinical settings. All samples were first sequenced using Illumina NGS whole-exome sequencing, and the pathogenic variant was known. The three patient samples involved different types of variants (shown in Fig. 6), including (1) a 10-base insertion in BCAP31; (2) a homozygous C > T SNP in SLURP1; and (3) two heterozygous C > T and T > C SNPs in UROC1. Using the standardized ECNano workflow, we obtained over 10 Gbp of base-called throughput with a single MinION flowcell and identified the target variants unambiguously. One of the three tested samples had a 10-based duplication prediction. Since the precision and sensitivity of INDEL-calling with ONT data is less promising than SNP-calling, the variant could still be called with high confidence. These test set results confirmed the robust discovery power of ECNano in actual clinical use.

**Fig. 6.**
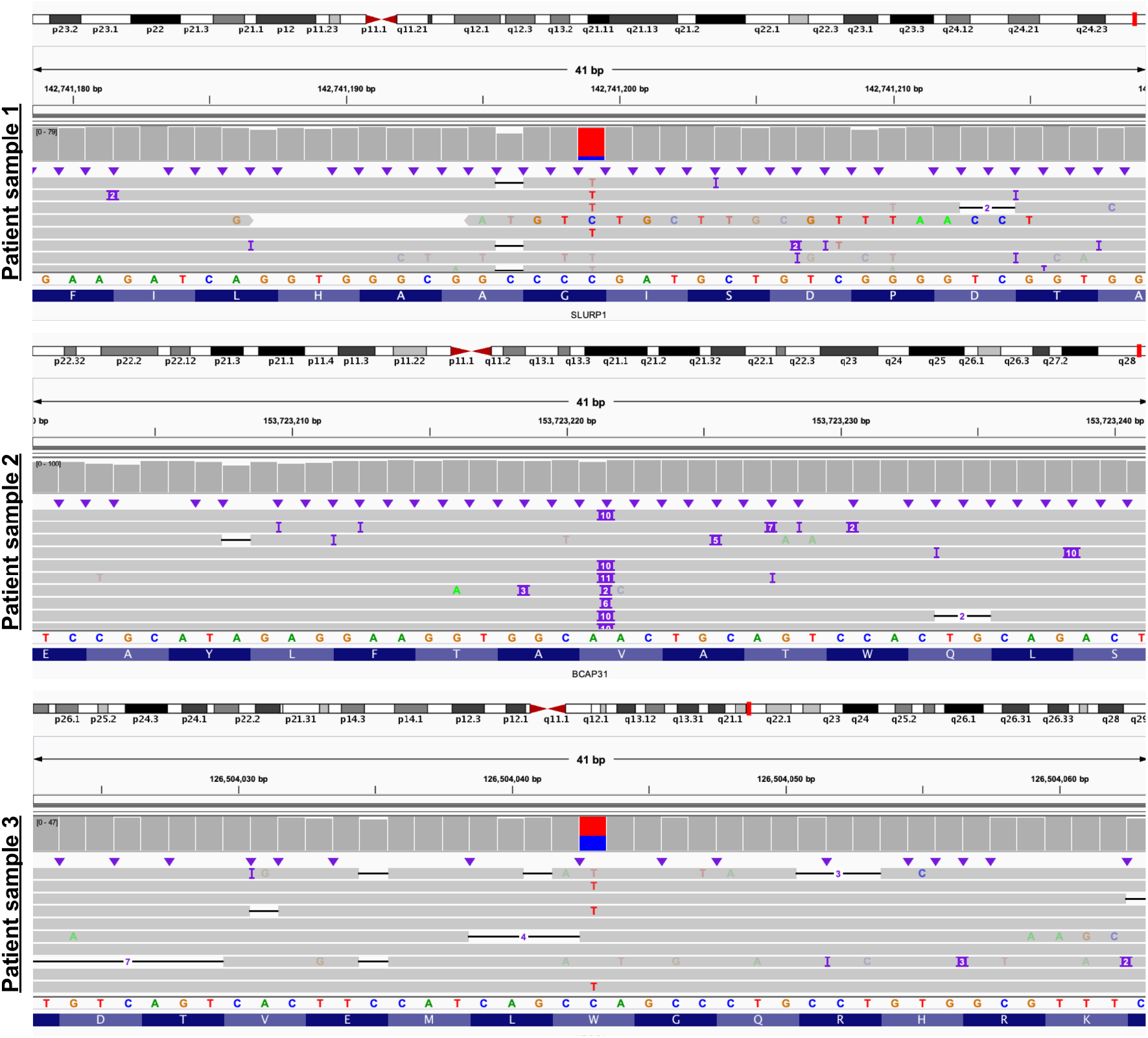
Middle position showing the target variant(s) in 3 patient samples and the alignment of the adjacent positions: patient sample 1 (top) with homozygous C > T SNP in *SLURP1*; patient sample 2 (middle) with 10-base insertion in *BCAP31*; and patient sample 3 (bottom) with heterozygous C > T SNP in *UROC1*.

## Discussion

ONT MinION sequencing facilitates long-read sequencing at lower cost and with short turnaround time by real-time base-calling. Initial checking for the variant was often observed within the first three hours of sequencing. Users can therefore flexibly decide their sequencing runtime based on current sequencing depth with reference to real-time read distribution. Medical applications of large panel target enrichment have high discovery power, providing comprehensive screening of all medically important genes. Compared with other multiple loci variant detection methods, such as DNA Array CGH (aCGH), MES has higher resolution and can potentially identify novel pathogenic positions via a population-wide association test and trio analysis. The sequencing cost, disk storage, and computational requirement of MES are also greatly reduced compared with whole genome sequencing (WGS), as the size of most-targeted sequencing panels shrinks below 5% of the whole genome [28]. In this study, we successfully integrated these two promising technologies together in a complete workflow, ECNano, for clinical use, with high-quality performance. We carefully selected the Agilent capture panel, which is a large panel, approximately 17Mbp in size, with medically important regions listed in three variant databases. Compared with other existing ONT target enrichment methods [29], ECNano uses hybrid-capture target enrichment for a large number of genes, is more stable and conventional, and is easier to incorporate into routine screening practices in clinical genetics labs. ECNano also provides a complementary bioinformatics workflow, Clair-ensemble, which is designed to process amplicon data. By adopting the original Clair, which performs rapid variant calling, Clair-ensemble improves the calling accuracy without a significant trade-off in the processing time. The whole ECNano workflow can be completed within 72 hours. This allows genetic diagnosis with ECNano to be rapid and precise.

There are areas yet to be explored in the application of the ECNano workflow, in addition to accurate variant calling. With such a high sequencing depth, intermediate size structural variance (SV of > 50bp up to 2,000bp) with a precise breakpoint within these exon regions or close by can be detected in principle [30]. The performance has been found to be poor using short reads [31]. Another potential application for ECNano sequencing is for variant phasing by region. Haplotype phasing allows more precise classification of the genetic configuration to better predict disease severity [32]. With ECNano long reads, more of these variants within individual target gene regions can be unambiguously phased.

## Conclusion

We presented a complete workflow, ECNano, for MinION sequencing of 4,800 clinically important genes and regions, including an optimized wet-lab protocol and a complementary bioinformatics pipeline, Clair-ensemble, for data processing and variant calling. In addition to the advantages of both hybridization-based target enrichment and long-read MinION sequencing, ECNano stably delivered high-quality results with a short turnaround time. Clair-ensemble allows accurate SNP calling by overcoming the ONT sequencing error and the uneven DoC among different captured regions. The long-read data has potential for further downstream analysis, such as variant phasing and intermediate size SV detection.

## List of abbreviations

aCGH: DNA Array CGH
DoC: Depth of coverage
MES: Medical exome sequencing
ONT: Oxford Nanopore Technologies
SNP: Single-nucleotide polymorphism
SNV: Single-nucleotide variants
SV: Structural variance
TGS: Third-generation sequencing
GS: Whole genome sequencing

## Declarations

### Ethics declarations

#### Ethics approval and consent to participate

The materials and protocols used in this study were reviewed and approved by the Human Research Ethics Committee (HREC) of HKU (reference no. EA210163). Informed consent was waived due to unidentified and anonymous data was used for new bioinformatic method development, the outcomes of which will not affect the standard of care and management of current patients. The experimental methods were in compliance with the Helsinki Declaration.

### Availability of data and materials

To ensure patient confidentiality, data containing potentially identifiable information was not shared. Raw fast5 data of ONT sequencing data obtained in our study is available from the corresponding author on reasonable request. The bioinformatics workflow, FASTQ of HG001 and HG002 datasets generated are available at GitHub (https://github.com/HKU-BAL/ECNano).

For the developed bioinformatics pipeline:

Project name: ECNano

Project home page: https://github.com/HKU-BAL/ECNano

Archived version: N/A

Operating system(s): Platform independent

Programming language: Python, C++, Shell

License: BSD 3-Clause License

Any restrictions to use by non-academics: No

## Authors’ contributions

HCML, RL and TWL conceptualized the study and revised the manuscript. AWSL and HCML designed the study and drafted the manuscript. AWSL performed the experiments. HML and IFML clinically examined the patients and collected clinical data and specimens. CLW, ZXZ and WWL developed the bioinformatics workflow. AWSL, HCML and ZXZ analyzed and interpreted the data. All co-authors have read the manuscript and agreed with its content. This manuscript was revised by all authors. All authors read and approved the final manuscript.

## Acknowledgements

We thank laboratory staff at the Clinical Genetic Service of the Department of Health (Hong Kong) for extracting DNA and performing NGS for the experiments.

## Supplementary data

**Additional file 1**.:

The performance of Clair-ensemble at target positions using two HG001 sequencing datasets and one HG002 sequencing dataset against the GRCh38 reference genome. The benchmarking results were compared against the original Clair settings with different models, LongShot and Medaka, using ECNano data. LongShot does not provide an INDEL calling function.

